# Motor planning of vertical arm movements in healthy older adults: does effort optimization persist with aging?

**DOI:** 10.1101/821314

**Authors:** Gabriel Poirier, Charalambos Papaxanthis, France Mourey, Jeremie Gaveau

## Abstract

Several sensorimotor modifications are known to occur with aging, possibly leading to adverse outcomes such as falls. Recently, some of those modifications have been proposed to emerge from motor planning deteriorations. Motor planning of vertical movements is thought to engage an internal model of gravity to anticipate its mechanical effects on the body-limbs and thus to genuinely produce movements that minimize muscle effort. This is supported, amongst other results, by direction-dependent kinematics where relative durations to peak accelerations and peak velocity are shorter for upward than for downward movements. The present study compares motor planning of fast and slow vertical arm reaching movements between eighteen young (24 ± 3 years old) and seventeen older adults (70 ± 5 years old). We found that older participants still exhibit directional asymmetries (i.e., differences between upward and downward movements), indicating that optimization processes during motor planning persist with healthy aging. However, the size of these differences was increased in older participants, indicating that gravity-related motor planning changes with age. We discuss this increase as the possible result of an overestimation of gravity torque or increased weight of the effort cost in the optimization process. Overall, these results support the hypothesis that feedforward processes and, more precisely, optimal motor planning, remain active with healthy aging.

## Introduction

Aging is associated with various sensorimotor modifications. Vision (Owsley 2011), vestibular function (Alberts et al. 2019), and proprioception (Goble et al. 2009) decline progressively. Walking and balance control are impaired (Laughton et al. 2003), and upper-limb arm movements become slower and more variable (Yan et al. 1998). These sensorimotor alterations lead to adverse effects such as falls, the most frequent cause of injuries in older adults (Grisso et al. 1990; World Health Organization. Ageing and Life Course Unit. 2008; Robinovitch et al. 2013). Because the number of older adults has fiercely increased in the past few decades, understanding the mechanisms responsible for age-related sensorimotor alterations is paramount.

Motor planning is a brain process specifying how a movement will be performed. Given a motor goal – e.g., grasping a cup of coffee – and a set of constraints – e.g., the gravitational pull on the arm and cup – this process selects the movement trajectory to be accomplished amongst a myriad of possible trajectories that would all attain the desired goal (Franklin and Wolpert 2011; Wong et al. 2015). For example, previous studies have shown that motor planning adapts reaching trajectories to changing inertial and gravitational constraints in young adults (Gaveau and Papaxanthis 2011; Vu et al. 2016a). There are pieces of evidence suggesting that aging alters motor planning (Kanekar and Aruin 2014; Kubicki et al. 2016; Casamento-Moran et al. 2017; Wunsch et al. 2017; Stöckel et al. 2017), and that altered motor planning may cause falls (Lord and Fitzpatrick 2001; Lyon and Day 2005; Robinovitch et al. 2013; Tisserand et al. 2016). Since fall is inherently linked to gravity, understanding how older adults adapt their motor planning to gravity is crucial.

Using arm reaching tasks, several studies have investigated gravity-related motor planning in young adults (Gentili et al. 2007; Le Seac’h and McIntyre 2007; Crevecoeur et al. 2009; Gaveau et al. 2011, 2014, 2016; Bringoux et al. 2012; Sciutti et al. 2012; Yamamoto and Kushiro 2014; Yamamoto et al. 2016, 2019; Hondzinski et al. 2016; Olesh et al. 2017). A consistent outcome is the existence of direction-dependent arm kinematics in normal gravity, which progressively disappears in microgravity (Papaxanthis et al. 2005; Gaveau et al. 2016). Direction-dependent kinematics is demonstrated by shorter time to peak acceleration and time to peak velocity for upward than for downward movements, i.e., more abrupt acceleration and softer deceleration. Model simulations explain these direction-dependent kinematics, and their progressive disappearance under microgravity conditions, as an optimization process that saves muscle effort (Berret et al. 2008; Crevecoeur et al. 2009; Gaveau et al. 2014, 2016). In young adults, direction-dependent kinematics has been proposed to represent the signature of a motor planning strategy that optimally integrates gravity torque to save muscle effort. Recent electromyographic (EMG) analyses further support this concept (Gaveau et al. 2019).

Here, relying on the above-described robust findings, we investigated the motor planning of vertical arm movements in young and healthy older adults. By comparing arm trajectories between upward and downward movements, we aimed to test whether older adults, like young ones, employ a motor planning strategy that optimally integrates gravity torque to save muscle effort.

## Methods

### Ethics and participants

This study was carried out following legal requirements and international norms (Declaration of Helsinki, 1964) and approved by the French National ethics committee (2019-A01558-49). Eighteen young (mean age = 24 ± 3 years; 14 males) and seventeen healthy older adults (mean age = 70 ± 5 years; 6 males) participated in this study after giving their written informed consent. All participants were right-handed, according to Edinburgh handedness inventory (Oldfield 1971), and did not present any neurological or musculoskeletal deficiency.

### Experimental design

Our experimental devices and protocol were similar to those of previous studies investigating motor planning of arm movements in the gravity field (Gentili et al. 2007; Gaveau et al. 2011, 2014, 2016). We investigated single-degree-of-freedom arm movements to manipulate the effects of gravity (with movement direction) while keeping the rest of the movement dynamics constant (inertial forces). Single degree-of-freedom movements allow to specifically investigating gravity-related motor strategies. Participants were comfortably seated on a chair with their trunk in a vertical position. The target system (a curved steel bar with 3 targets fixed on it) was vertically aligned and placed in front of the participants’ right arm at a distance equal to the length of their fully extended arm plus two centimeters (Figure 1A). We horizontally aligned the central target (starting target) with the participants’ shoulder, while positioning the other two targets to imply a 30° upwards or a 30° downwards shoulder rotation. Starting from the central target, participants performed single-degree-of-freedom (rotation around the shoulder joint) upward and downward arm movements, as accurately as possible at two speeds. Previous work revealed that movement speed significantly influences the temporal organization of vertical arm movements (Gaveau and Papaxanthis 2011). Therefore, to obtain movements of comparable durations between conditions and groups, automatic online analyses of movement duration were performed, and the results were used to provide verbal feedback to the participants. This feedback aimed at driving participants to perform reaching movements lasting about 350ms (called fast speed below) and 550ms (called slow speed below). Fast and slow trials were performed in a random block design, and upward and downward movements were randomly ordered within each block (12 trials × 4 conditions; 48 trials overall for each participant). Participants performed a few practice trials before the beginning of the experiment (~ 10 practice trials per experimental condition). At the beginning of each trial, they were invited to point at the central target (starting position, see Fig. 1A). After a short period (~2s), the experimenter verbally indicated the target (upward or downward) to reach. Participants were then allowed to carry out the movement without any constraint on their reaction time and to maintain their final finger position for about 2 seconds until a verbal signal informed them to relax their arm. In order to avoid muscle fatigue, we separated each trial by a short rest period (~15s) and both blocks by a five minutes rest period.

**Figure 1:**
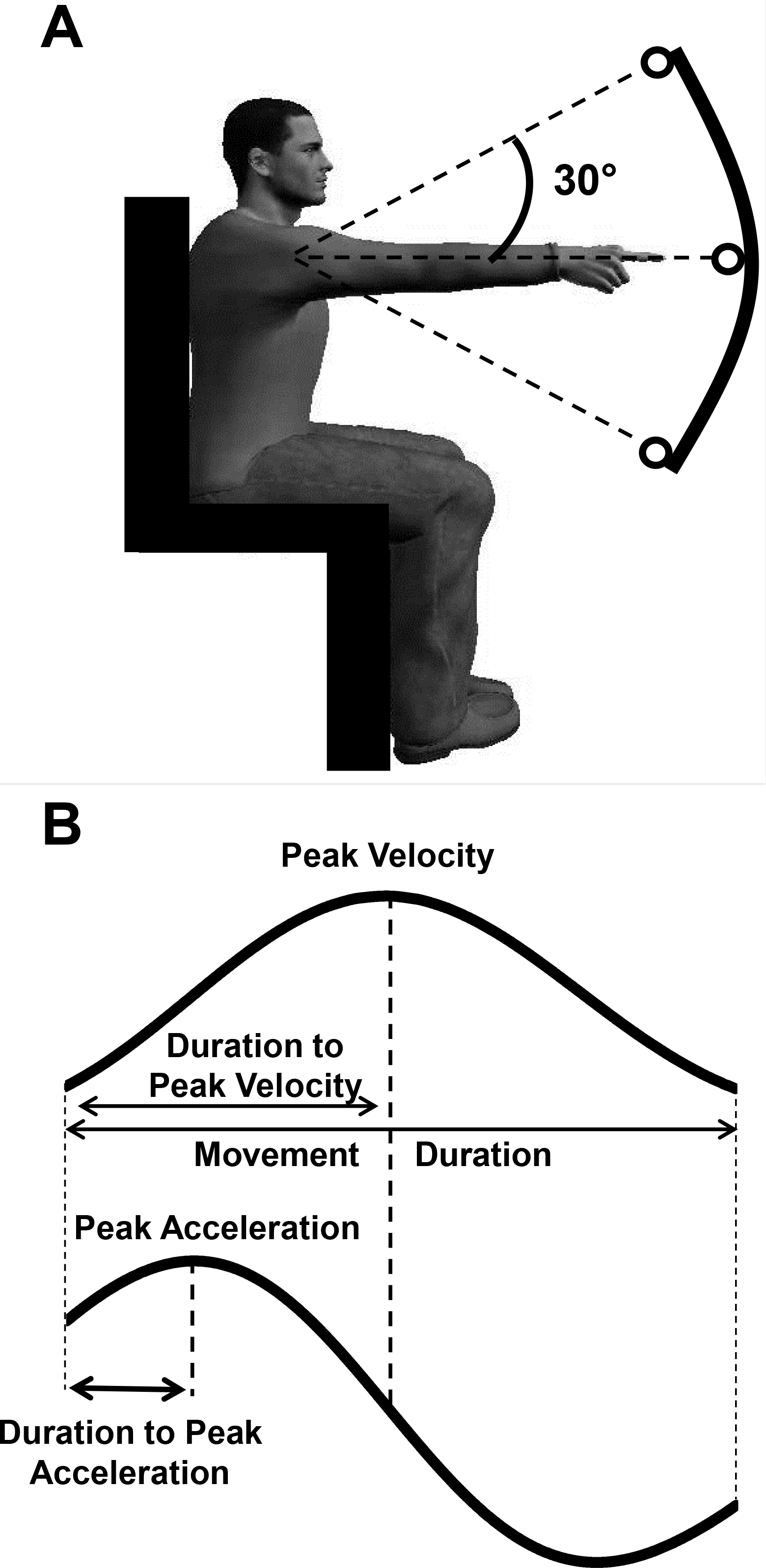
**A.** Experimental setup and participant’s starting position. From this starting position, participants pointed to the upward or the downward target on separated trials. **B.** Illustration of the parameters computed on the velocity and acceleration profiles.

We placed five reflective markers on the participants’ right shoulder (acromion), arm (middle of the humeral bone), elbow (lateral epicondyle), wrist (cubitus styloid process), and finger (nail of the index). We recorded markers’ position with an optoelectronic motion capture system (VICON system) that consisted of six cameras (sampling frequency: 100Hz; precision < 0.5mm).

### Data analysis

Data were processed using custom programs written in Matlab (Mathworks, Natick, MA). The tridimensional position (X, Y, Z) of markers was low-pass filtered with a digital third-order Butterworth filter (zero-phase distortion) at a cut-off frequency of 10Hz. We derived these signals in order to compute velocity and acceleration profiles for each movement. Movement start and end was defined as the moment when finger velocity respectively went above or fell below 10% of the peak velocity value (Gaveau and Papaxanthis 2011; Gaveau et al. 2014, 2016). Movement duration and amplitude were computed based on movement start and end.

Following results from previous studies, we quantified the temporal organization of each movement with two temporal parameters that are theoretically independent of the movement duration (see Figure 1B and Gaveau *et al.*, 2011, 2014); i) the relative duration to peak acceleration (rD-PA), defined as the duration to peak divided by movement duration; ii) the relative duration to peak velocity (rD-PV), defined as the duration to peak velocity divided by movement duration.

### Statistical analysis

We used STATISTICA 10 (StatSoft. Inc.) to perform all statistical analyses. All variables showed a normal distribution (Kolgomorov-Smirnov test). We carried out repeated measures variance analysis (three-way ANOVA) with 3 factors: *age* (Young vs Older), *speed* (Fast and Slow), and *direction* (Upward vs. Downward). We used *Scheffé tests* for post hoc comparisons. The level of significance was set at p<0.05 for all analyses.

## Results

Participants accurately reached targets (average movement amplitude: 28.8 ± 1.1 ° for young and 28.7 ± 2.1° for older participants, see Table 1 for all values) with smoothed movement exhibiting single-peaked and bell-shaped velocity profiles (Kelso et al. 1979). Figure 2 qualitatively illustrates the mean position, velocity, and acceleration profiles recorded for each group and direction at the fast speed.

**Table 1:** Kinematic parameters for each condition. Mean (±SD) movement duration (in ms), amplitude (in °), relative duration to peak acceleration (rD-PA), and relative duration to peak velocity (rD-PV) are presented for each experimental condition.

|  | Fast |  |  |  | Slow |  |  |  |
| --- | --- | --- | --- | --- | --- | --- | --- | --- |
|  | Young |  | Older |  | Young |  | Older |  |
|  | Up | Down | Up | Down | Up | Down | Up | Down |
| **Movement Duration (ms)** | 328 ± 31 | 330 ± 30 | 349 ± 52 | 347 ± 51 | 570 ± 147 | 573 ± 145 | 581 ± 115 | 578 ± 135 |
| **Amplitude (°)** | 29.6 ± 1.1 | 29.0 ± 1.1 | 29.2 ± 1.7 | 30.3 ± 2.5 | 28.4 ± 1.4 | 28.2 ± 1.1 | 27.6 ± 2.1 | 27.9 ± 2.0 |
| **rD-PA** | 0.23 ± 0.01 | 0.24 ± 0.01 | 0.20 ± 0.03 | 0.23 ± 0.02 | 0.19 ± 0.04 | 0.21 ± 0.03 | 0.18 ± 0.03 | 0.22 ± 0.03 |
| **rD-PV** | 0.48 ± 0.02 | 0.49 ± 0.02 | 0.44 ± 0.04 | 0.47 ± 0.03 | 0.46 ± 0.04 | 0.47 ± 0.04 | 0.43 ± 0.04 | 0.47 ± 0.04 |

**Figure 2:**
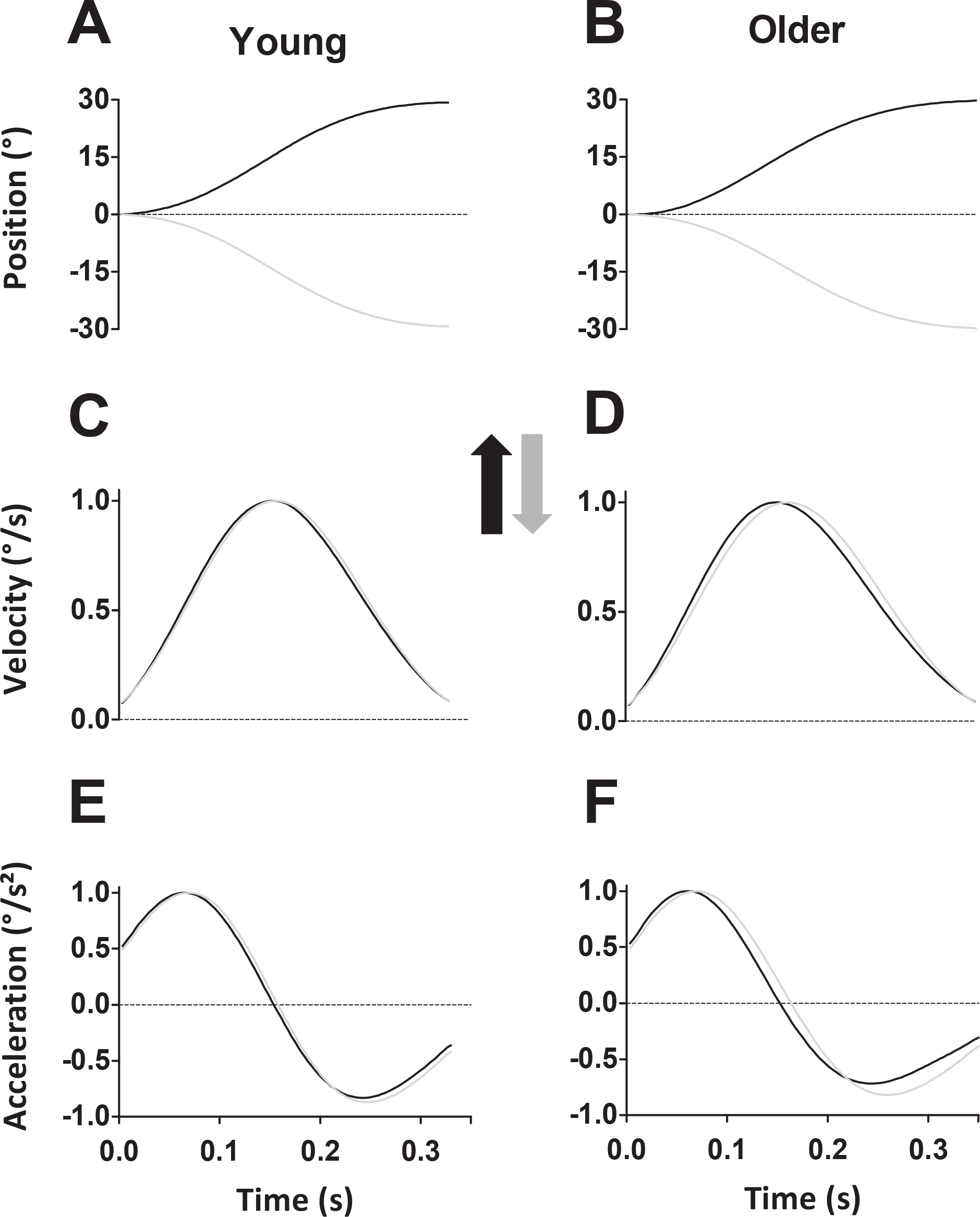
Mean position (**A-B**), velocity (**C-D**), and acceleration (**E-F**) profiles for each group and direction at the fast speed. Black traces represent upward movements, and grey traces represent downward movements. Left panels display young participants’ data, while right panels display older participants’ data.

### Movement Duration

Because movement duration was previously shown to influence the temporal organization of arm movements (Gaveau and Papaxanthis 2011), to ensure that it was similar between conditions, we performed the same statistical analysis on movement duration as on the other parameters. As required by task constraints, movement duration significantly varied with speed instructions (*speed* effect: F = 105.18; p<0.001; η_p_^2^ = 0.76). Average movement durations were respectively 340 ± 40 ms and 575 ± 120 ms for fast and slow movements (Table 1). However, neither *age* (F = 0.27; p = 0.60; η_p_^2^ = 0.008) nor *direction* (F = 0.001; p = 0.97; η_p_^2^<0.001) effects were observed. Similarly, all interaction effects (*age* × *direction*, F = 0.39; p = 0.53; η_p_^2^ = 0.01; *speed* × *age*, F = 0.06; p = 0.81; η_p_^2^ = 0.001; *speed* × *direction*, F = 0.016; η_p_^2^<0.001; p = 0.90; *age* × *speed* × *direction*, F = 0.10; p = 0.92; η_p_^2^<0.001) were non-significant. These results show that movement duration was similar between groups and directions within each speed instruction.

### Relative Duration to Peak Acceleration (rD-PA)

Figure 3A and B displays rD-PA for each group, speed, and direction (see also Table 1). rD-PA was smaller for upward than for downward movements (*direction* effect, F = 67.95; p<0.001; η_p_^2^ = 0.67). Comparison between Figure 3A and B also reveals that rD-PA was smaller for slow than for fast movements (*speed* effect, F = 40.64; p<0.001; η_p_^2^ = 0.55). Of particular interest for the present study is the fact that the difference between upward and downward movements was larger in the older than in the young adults at both speeds. The ANOVA yielded a significant *age* × *direction* interaction effect (F = 4.41; p = 0.04; η_p_^2^ = 0.12) but no *age* × *speed* × *direction* interaction (F = 0.003; p = 0.95; η_p_^2^<0.001). Box-plot in Figure 3C presents the ratio of directional difference ((Down-Up)/Down) to further illustrate the effect of age and speed on direction-dependent kinematics.

**Figure 3:**
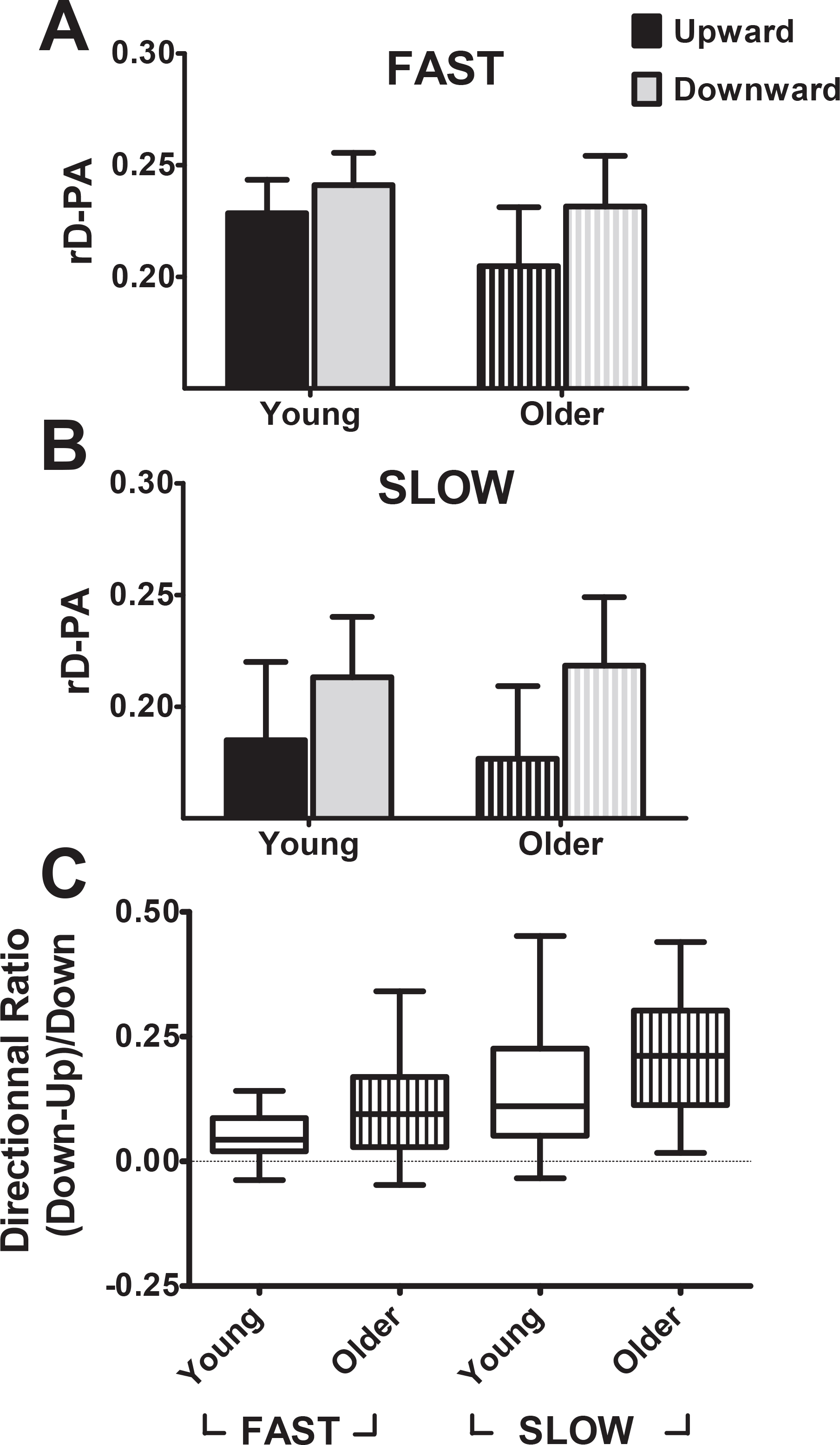
**A.** Mean (± SD) relative duration to peak acceleration (rD-PA) for upward (black) and downward (grey) fast movements. Solid bars represent young, and striped bars represent older participants. **B.** Mean (± SD) relative duration to peak acceleration (rD-PA) for upward (black) and downward (grey) slow movements. Solid bars represent young, and striped bars represent older participants. **C.** Box plots show directional ratio (computed for each participant as: ((rD-PA_Down_ - rD-PA_Up_)/rD-PA_Down_) for young (empty boxes) and older (striped boxes) participants for fast and slow movements. Whiskers represent 95% confidence interval.

### Relative Duration to Peak Velocity (rD-PV)

As for rD-PA, the relative duration to peak velocity was shorter for upward compared to downward movements (F = 20.21; p<0.001; η_p_^2^ = 0.38) and for slow compared to fast movements (F = 4.98; p = 0.03; η_p_^2^ = 0.13; see Table 1). Also, the *age* × *direction* interaction yielded a significant effect (F = 9.21; p = 0.005; η_p_^2^ = 0.22), further supporting the increased directional asymmetry in older compared to young adults. As for the rDPA, the *age* × *speed* × *direction* interaction did not reach significance (F = 0.02; p = 0.90; η_p_^2^ <0.001).

## Discussion

We investigated the motor planning of single-degree-of-freedom vertical arm movements in young and older adults. Precisely, we compared directional asymmetries (the difference between upwards and downwards) on relative durations to peak acceleration and peak velocity. Our results show increased directional asymmetries on both parameters in older participants compared to young ones.

In young adults, several studies have shown that single-degree-of-freedom arm movements exhibit directional asymmetries in the vertical plane. These asymmetries: i) were consistently reported as a more abrupt acceleration phase for upward than for downward movements; ii) do not exist in the horizontal plane (Gentili et al. 2007; Le Seac’h and McIntyre 2007); iii) progressively disappear in the vertical plane when participants are exposed to microgravity (Papaxanthis et al. 2005; Gaveau et al. 2016); iv) were observed very early in the movements (before 70ms after movement start (Gaveau and Papaxanthis 2011; Gaveau et al. 2014); v) were explained by optimal control model simulations and EMG analyses as a motor strategy that minimizes muscle effort (Berret et al. 2008; Crevecoeur et al. 2009; Gaveau et al. 2014, 2016, 2019). Altogether, in young adults, previous results strongly suggest that direction-dependent kinematics represent the signature of a motor planning strategy that optimally integrates gravity torque to save muscle effort. The present results reveal that older adults also produce directional asymmetries in the vertical plane. Most importantly, these asymmetries have the same sign as those of young adults – a more abrupt acceleration for upward than for downward movements – and they emerge early in the movement – rD-PA was already different, which is about 70ms after movement onset at the fastest speed –. First, this reveals that older adults, similar to young adults, plan arm movements that are direction-dependent in the gravity field. Second, the present results suggest that optimal motor planning, and more specifically, the minimization of muscle effort, continue with healthy aging. Indeed, although the size of the directional asymmetry was shown larger in older compared to young adults, previous studies have demonstrated that the simple existence of the directional asymmetry – independently of its size – is sufficient to reveal an effort-related optimization process (Berret et al. 2008; Gaveau et al. 2016). Hereafter, building on recent experimental and modeling results, we discuss the increased direction-dependence of arm kinematics in older adults.

Studying motor planning within the optimal control framework, previous works explained human movement as the minimization of composite cost functions (for a recent review, see Berret et al. 2019). For example, minimizing a trade-off between effort and smoothness allowed to explain how reach endpoint is selected during multi-articular arm movement tasks (Berret et al. 2011; Vu et al. 2016b). Using an effort and smoothness composite cost function, the model simulations from Gaveau et al. (2016) also demonstrated that increasing the weight of the effort cost produces an increase in directional asymmetries. Thus, the increased importance of effort during motor planning could explain the increased directional asymmetry observed in older adults. Indeed, aging is associated with a loss of muscle mass and strength (for a review, see Mitchell et al. 2012) that likely results in an increased perception of effort for daily-life tasks (McCloskey et al. 1974; Hess et al. 2016; Pageaux and Lepers 2016).

Experimental and modeling results have also demonstrated that the size of the directional asymmetry is proportional to the gravity torque (Gaveau et al. 2014, 2016). Increasing the gravity torque either by moving body limbs of increasing mass (i.e., upper arm, forearm, and wrist) or by adding an external load on the limb both led to increased directional asymmetries in the vertical plane (Gaveau et al. 2014). Progressively varying movement direction relative to gravity caused a progressive variation of the temporal organization of arm movement such that arm kinematics and gravity torque linearly correlated with each other (Gaveau et al. 2016). Last, during exposure to microgravity, model simulations explained the progressive disappearance of directional asymmetries as the progressive decrease of the gravity value, i.e., a recalibration of the internal gravity model (Gaveau et al. 2016). Thus, the anticipation of an increased gravity torque could explain the increased directional asymmetry observed in older adults. Experiments on object weight perception in older adults have provided support for such an overestimation of gravity torque (Parikh and Cole 2012; Holmin and Norman 2012). Assuming that motor control is the result of two parallel processes, a forward model that produces accurate motor output and an optimal controller selecting the trajectories that minimize some hidden motor costs (Izawa et al. 2008), over-estimating gravity torque would lead to new optimal trajectories without impeding movement accuracy (Gaveau et al. 2011).

Sensory information about gravity is known to influence motor planning (Bringoux et al. 2012; Rousseau et al. 2016). With aging, several studies have provided evidence for sensory deterioration in visual (see Saftari and Kwon (2018) for a review), proprioceptive (Goble et al. 2009), and vestibular systems (Alberts et al. 2019), as well as their multisensory weighting (de Dieuleveult et al. 2017). The perception of gravity vertical also deteriorates with aging (Kobayashi et al. 2002; Baccini et al. 2014). One may, therefore, wonder whether an alternative hypothesis for increased directional asymmetry in older adults could be the elderly’s’ failure to measure gravity torque. In other words, one may expect older adults to apply the same motor plan for vertical and horizontal movements. This premise is implausible as it would produce either no directional asymmetry (Gentili et al. 2007; Le Seac’h and McIntyre 2007) – should the forward model be accurate – or extensive end-point errors and directional asymmetries with an opposite sign – should the forward model be inaccurate –. We observed none of those effects in the present study. Conversely, it is crucial to underline that we observed similar results for rD-PA and rD-PV. Also, we observed no interaction and small effect sizes of *age* × *speed* × *direction*, indicating that speed did not influence the age effect on directional asymmetries. Since rD-PV happens late in the movement, especially at slow speed (about 250ms after onset), our results suggest that no online feedback-driven correction was implemented to address hypothesized motor planning errors.

The present results add to the existing literature suggesting that motor planning is modified with aging (Kanekar and Aruin 2014; Kubicki et al. 2016; Casamento-Moran et al. 2017; Wunsch et al. 2017; Stöckel et al. 2017). Neuroscientists have first interpreted motor planning modifications as a deterioration of feedforward processes (proactive strategies) that urges older adults to favor feedback processes (reactive strategies). The most potent experimental support for this hypothesis is the general observation that movements become slower with aging (Buckles 1993; Yan et al. 1998). However, older adults are not always slower than younger adults. Asking participants to reach, grasp and lift an object at their own pace, Hoellinger et al. (2017) found that older adults moved faster than young adults. Comparing their empirical results to model simulations, the authors suggested that older adults planned stronger feedforward forces in order to favor feedforward mechanisms over feedback ones. This hypothesis is rooted in results showing that the sensory system becomes noisier with aging (Desmedt and Cheron 1980; Doherty et al. 1994; Stevens and Choo 1996). Thus, to compensate for their noisy unreliable sensory system, older adults may favor feedforward over feedback mechanisms. This hypothesis is well-supported by recent results showing that sensory attenuation, a well-studied feedforward mechanism (Blakemore et al. 1998; Shergill et al. 2005; Pareés et al. 2014), increases with aging (Wolpe et al. 2016). Results of neurobiological studies suggesting that increased brain activations compensate for neuro-behavioral deficits to preserve motor performance in older adults could also support the hypothesis of increased reliance on feedforward mechanisms (Cabeza 2002; Berchicci et al. 2012).

There are pieces of evidence suggesting that feedforward mechanisms remain functional with healthy aging and may even compensate for unreliable feedback mechanisms (Boisgontier and Nougier 2013; Wolpe et al. 2016; Helsen et al. 2016; Hoellinger et al. 2017; Vandevoorde and Orban de Xivry 2019). The result that older adults still plan direction-dependent arm movements in the gravity field further supports and extends this hypothesis. The present study also reveals that aging exacerbates the direction-dependence of arm kinematics. This result constitutes the first insight into the effect of aging on gravity-related motor planning, whose better understanding may benefit the prevention and rehabilitation of falls and fallers in older adults. Indeed, although we investigated single degree of freedom arm movements to isolate gravity effects here, it is essential to mention that directional asymmetries are also documented for multi-articular arm reaching, reaching to grasp, grasping, hand drawing, and whole-body sit-to-stand / stand-to-sit movements (Papaxanthis et al. 1998, 2003, 2005; Yamamoto and Kushiro 2014). Future studies should attempt to extend the present findings to whole-body movement tasks and to disentangle sensory from motor planning modifications in older adults.

## Acknowledgments

This work was supported by the Institut National de la Santé et de la Recherche Médicale (INSERM) and by the Agence National de la Recherche (ANR, project MOTION ANR-14-CE30-0007-01). We thank Cyril Sirandré and Yves Ballay for their technical support.

